# The kinesin-14, Ncd, drives a right-handed, helical motion of antiparallel microtubules around each other

**DOI:** 10.1101/757914

**Authors:** Aniruddha Mitra, Rojapriyadharshini Gandhimathi, Felix Ruhnow, Roman Renger, Stefan Diez

## Abstract

Within the mitotic spindle, several kinesin motors crosslink and slide microtubules. While some of them (e.g. kinesin-5, kinesin-8 and kinesin-14) have been shown to exhibit sideways components in their step cycles, the impact of the resulting off-axis power strokes on motility and force generation in the spindle has not been investigated so far. Here, we develop and utilize a novel three-dimensional *in vitro* motility assay to explore the kinesin-14, Ncd, driven sliding of crosslinked, fluorescently-labeled microtubules. We find that free microtubules, sliding in an antiparallel orientation on microtubules suspended between nanofabricated ridges, not only rotate around their own axis but also move around the suspended microtubules with right-handed helical trajectories. In contrast, microtubules crosslinked in parallel orientation are static with neither longitudinal nor helical motion. Further, our technique allows us to measure the *in situ* spatial extension of the motors between the crosslinked microtubules to be about 20 nm. We argue that the capability of microtubule-crosslinking kinesins to cause helical motion of microtubules around each other allows for flexible filament organization, roadblock circumvention and torque generation in the mitotic spindle.

## Introduction

The mitotic spindle is a complex subcellular machinery that segregates chromosomes during eukaryotic cell division. Several *in vivo* and *in vitro* studies provide us with a consolidated twodimensional (2D) model, detailing the mechanisms employed by dynamic microtubules, kinesins and microtubule-associated proteins to assemble, maintain and disassemble the spindle in order to coordinate chromosome segregation^1,2^. However, the spindle is a threedimensional (3D) structure and interesting mechanical details may emerge in the third dimension, which are not just intuitive extensions of the 2D model. For instance, a recent study suggests that the mitotic spindle is twisted with a distinct right-handed chirality^3,4^.

Interestingly several mitotic kinesins, like kinesin-5^5^, kinesin-8^6,7^ and kinesin-14^8,9^, have been shown to exhibit off-axis components in their stepping behavior, i.e. they do not move strictly parallel to the longitudinal microtubule axis. These mitotic kinesins are microtubule crosslinkers involved in connecting and sliding microtubules ^10–12^. To address if their off-axis (i.e. axial) motion translates to crosslinked microtubules, we explore the 3D motion of microtubules propelled along each other by the kinesin-14, Ncd. Ncd is a non-processive, minus-end directed motor possessing an additional diffusive microtubule-interacting site in its N-terminal tail domain that enables the motor to crosslink and slide anti-parallel microtubules^10^. Specifically, we suspend long microtubules on nano-fabricated ridges^7,13^ and track the 3D motion of short microtubules along the long ones. We discover that anti-parallel microtubules do both, rotate around their own axes and helically move around each other, in a right-handed manner.

## Results

To study the uninhibited sliding motion of crosslinked microtubules driven by *Drosophila melanogaster* kinesin-14, Ncd, motors, we suspended parts of long ‘template’ microtubules such that short crosslinked ‘transport’ microtubules could access the entire lattice of the suspended template microtubules. As illustrated in Fig. 1A, rhodamine-labeled template microtubules were tautly fixed on optically-transparent polymer ridges (370 nm high and 2 or 5 μm wide) separated by 10 μm wide valleys via anti-rhodamine antibodies, as developed in previous studies^7,13^. Atto647n-labeled transport microtubules were crosslinked to the template microtubules via full length GFP-Ncd (hereafter referred to as Ncd) in ADP. After the application of ATP, antiparallel transport microtubules started to slide along the template microtubules while parallel transport microtubules remained locked in their position. In order to observe the position of the transport microtubules with respect to the template microtubules, the microtubule positions were tracked in 2D individually in the rhodamine and Atto 647n fluorescence channels (using FIESTA^14^) and the distances of the center points of the transport microtubules from the tracked center lines of the template microtubules, referred to as the sideways distance, were obtained (Fig. 1B). Since the imaging set-up does not provide information about the third dimension, the sideways distances provide the xy-projected distances between the crosslinked microtubules, which are maximum when the transport microtubules are at the extreme left (or right) of the template microtubules and zero when the transport microtubules are on top (or bottom) of the template microtubules.

**Figure 1:**
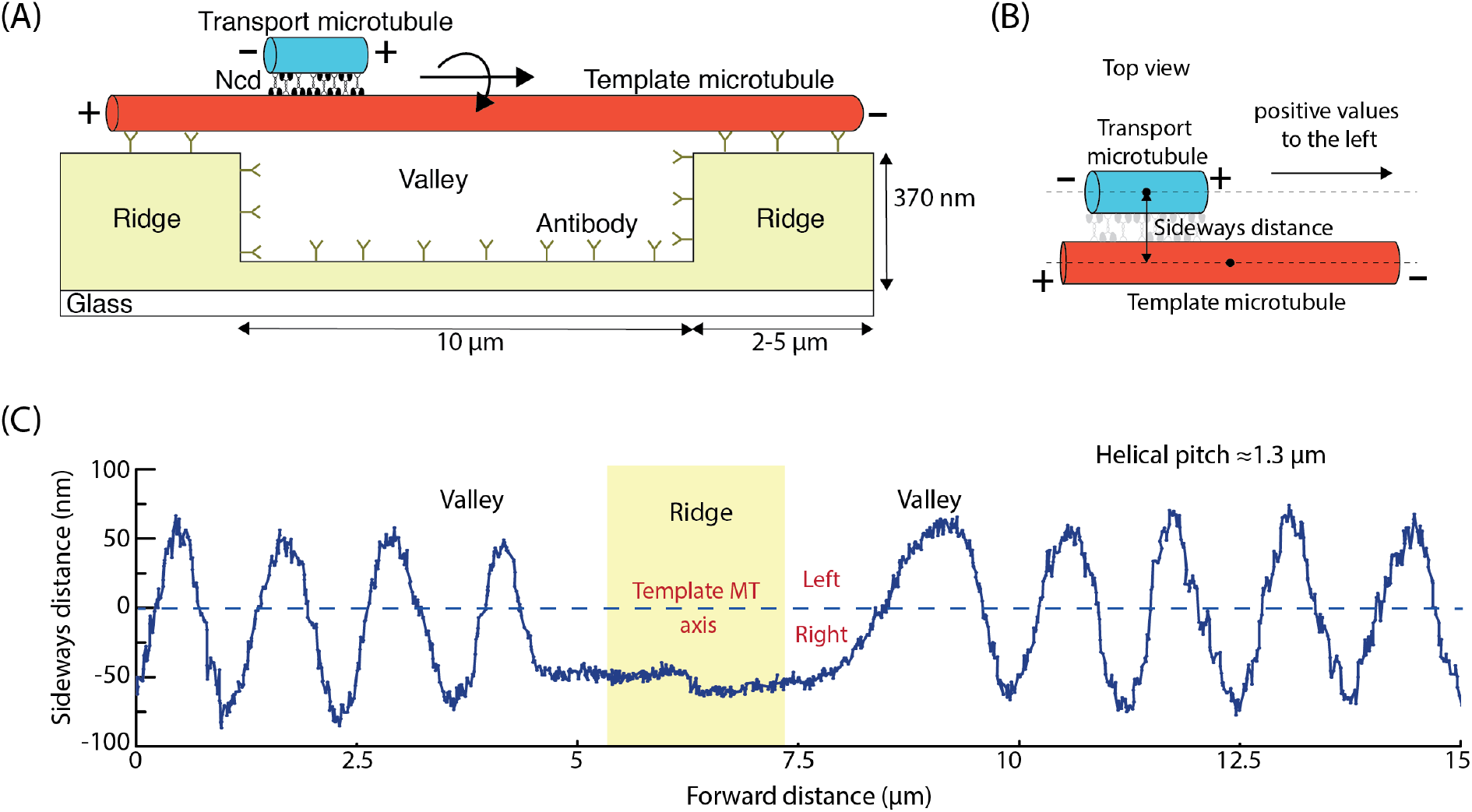
3D sliding of transport microtubules driven by Ncd on suspended template microtubules. **(A)** Schematic representation of a suspended, rhodamine-labeled, template microtubule immobilized on optically-transparent polymer ridges by anti-rhodamine antibodies. Atto647n-labeled transport microtubules are capable of freely accessing the 3D lattice of the template microtubule between the ridges (in the region referred to as valley), as they slide along them driven by GFP-labeled Ncd motors. **(B)** The perpendicular distance of the center point of the transport microtubule from the tracked center line of the template microtubule provides the sideways distance. Positive values are arbitrarily assigned to positions on the left. **(C)** Sideways distance for an example antiparallel transport microtubule as it slides along a template microtubule over two valley regions (see also Supplementary Movie 1). In both valley regions, the 1.8 μm long transport microtubule exhibited a helical motion around the suspended template microtubule, with a helical pitch of 1.3 ± 0.1 μm (N = 7 rotations, mean ± SD). At the ridge, where the template microtubule is surface-immobilized, the transport microtubule remained on the right-hand side of the template microtubule, indicating that the direction of helical motion is right-handed.

To interpret the 3D motion of antiparallel transport microtubules the sideways distance was plotted with respect to the distance travelled along the longitudinal axis of the template microtubules. Fig. 1C (and Supplementary Movie 1) shows an exemplary sliding event at 4 nM Ncd (in solution), where a transport microtubule moved along a template microtubule suspended over two valleys with a 2 μm wide ridge in between. In the valley regions, the transport microtubule followed a helical trajectory with a pitch of 1.3 ± 0.1 μm (mean ± S.D.; N = 7 complete rotations). When the transport microtubule encountered the ridge, its helical motion was blocked without significant impediment to the longitudinal motion, and it stayed at the right-hand side of the template microtubule. The transport microtubule only resumed its helical motion when the entire microtubule (1.8 μm long) had left the ridge. This indicates that surface immobilization of the template microtubule on the ridge hindered the helical motion of the transport microtubule. Since the helical motion of the transport microtubule on the ridge was locked on the right-hand side of the template microtubule, we infer that the helical motion is right-handed (or clockwise in the direction of motion).

We analyzed 125 microtubule-microtubule interaction events on suspended template microtubules at a Ncd concentration of 4 nM in solution. *Antiparallel transport microtubules* exhibited robust helical trajectories over the valleys along suspended template microtubules (see 30 example events in Fig. 2A and all 94 rotating events in Supplementary Fig. 1A). Interestingly, while the helical pitches within individual events were relatively constant, there was a sizable variation in helical pitches between events (ranging from 0.5 μm to 3 μm as seen in Supplementary Fig. 1B; median helical pitch 1.6 ± 0.2 μm; see methods for analysis of distributions with error estimation) often even on the same template microtubule. There was no evident correlation between the helical pitches and the sliding velocities (Supplementary Fig. 1C). Also, the helical pitches were not correlated with the lengths of the transport microtubules, though shorter microtubules exhibited larger variations in helical pitches (Supplementary Fig. 1D). When sliding on the ridge-immobilized parts of the template microtubules, transport microtubules were almost always on the right-hand side of the template microtubules (Fig. 2B), with a mean sideways distance of - 41.6 ± 0.4 nm (N = 63). This clearly indicates that the helical motion induced by Ncd is right-handed. Only in a couple of events the transport microtubules move around the surface-immobilized template microtubules, presumably at locations of statistically low densities of tubulin antibodies. These observations are consistent with 2D sliding motility assays performed on unstructured glass coverslips (Supplementary Fig. 2A, example event in Supplementary Movie 2). In this assay geometry, transport microtubules moved either along the right-hand side of the template microtubules or flipped around the template microtubules with aperiodic helical motion (Supplementary Fig. 2B). No clear dependence on motor density was observed (experiments performed at Ncd concentrations of 0.4 nM, 4 nM and 40 nM). The mean sideways distance was - 44.6 ± 0.1 nm (N = 98) and upon removing the flip events became - 45.5 ± 0.1 nm (N = 83). At lower Ncd concentration (0.02 nM), the sideways motion of transport microtubules became erratic. However, there was still a slight bias to the right-hand side with a mean sideways distance of −26.8 ± 0.7 nm (N = 24). *Parallel transport microtubules* were found to be locked, both in their longitudinal and helical motion, and did not exhibit any preference for which side of the template microtubule they were bound to (mean sideways distance 13.7 ± 1.0 nm; N = 30, Fig. 2C). Rather, the distribution of sideways distances was bimodal, consistent with the distribution of microtubule protofilaments projected in 2D and an equal probability of transport microtubules interacting randomly with any protofilament of the template microtubules, providing peaks at about ± 45 nm. Similar to our observations on suspended microtubules, parallel microtubules on surface-immobilized template microtubules were also locked, though with a minor bias to the right (mean sideways distance of −34.8 ± 0.6 nm; N = 52; Supplementary Fig. 2C). In summary, transport microtubules that were antiparallel to the template microtubules showed robust right-handed helical motion on suspended template microtubules with helical pitches of 0.5 - 3 μm and their helical motion was hindered on surface-immobilized template microtubules. In contrast, parallel transport microtubules appeared to be locked in the longitudinal as well as the axial direction.

**Figure 2:**
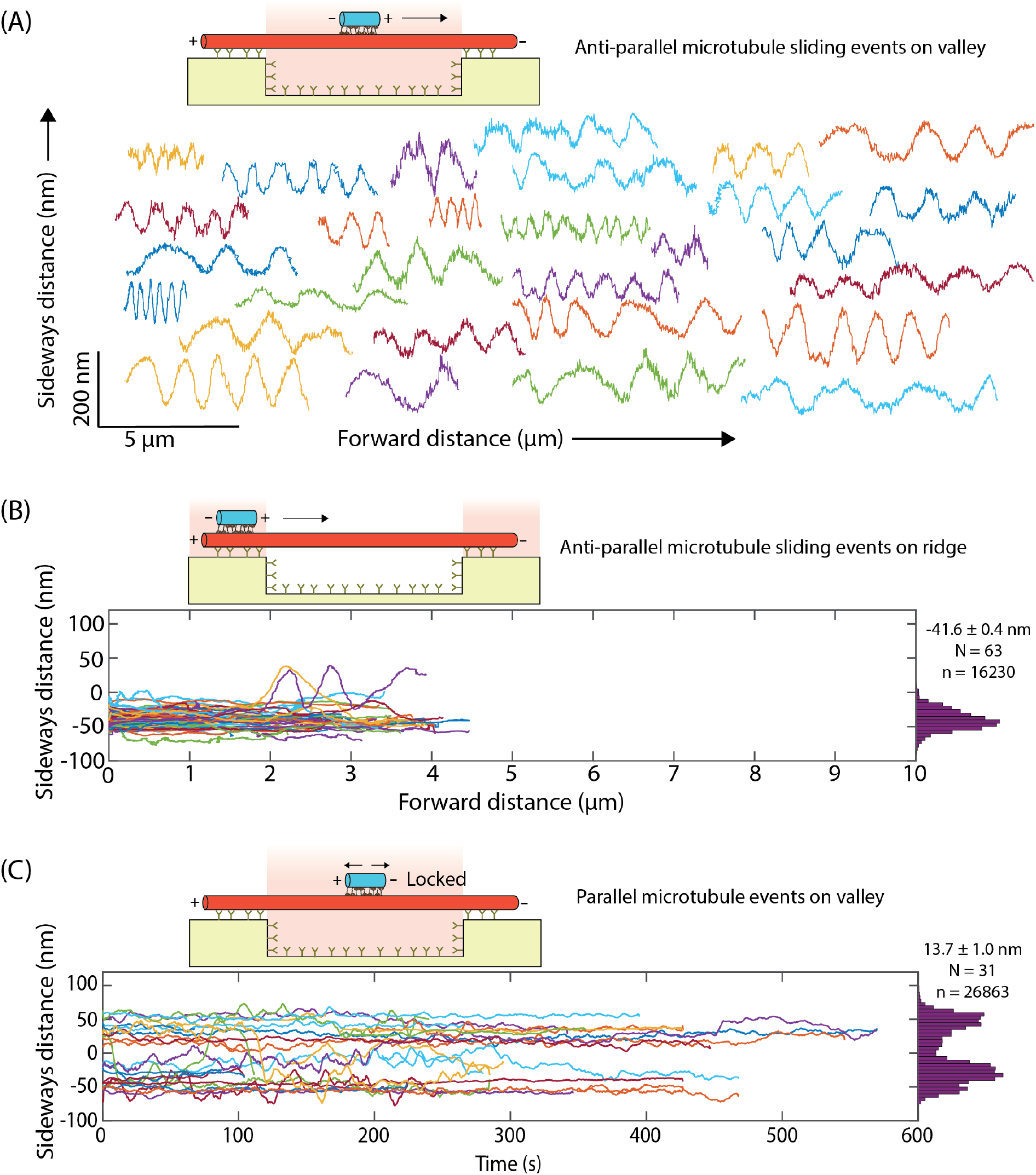
Trajectories of transport microtubules antiparallel and parallel to the template microtubules. **(A)** Sideways distance of 30 example antiparallel transport microtubule (out of 94 events; all events shown in Supplementary Figure S1A) driven by Ncd along the suspended parts of template microtubules. Each transport microtubule exhibited a robust helical motion with relatively constant helical pitch. Between various transport microtubules, helical pitches varied significantly between 0.5 μm and 3 μm. **(B)** Sideways distance of 63 antiparallel transport microtubules along the ridge-immobilized parts of the template microtubules. Most transport microtubules remained on the right-hand side of the template microtubule while a couple of microtubules managed to squeeze under the template microtubules to perform complete rotations. The average sideways distance was −41.6 ± 0.4 nm (N = 63 events; n = 16230 data points; see methods for analysis of distributions with error estimation). **(C)** Sideways distance (plotted with respect to time) of 30 parallel transport microtubules along the suspended parts of template microtubules. The average sideways distance was 13.7 ± 1.0 nm (N = 30 events; n = 26863 data points). All events showed very little (< 300 nm) or no motion in the longitudinal direction. Transport microtubules were additionally locked in the axial direction with no preference for either side of the template microtubule. Only events on template microtubules having one or more antiparallel microtubule sliding events were chosen, in order to define whether the transport microtubule is on the left or right-hand site of the template microtubule. In (B) and (C) the smoothened trajectories are plotted (rolling frame averaged over 20 frames).

Intriguingly, our technique additionally allows us to gauge the *in situ* spatial extension of the motors between sliding microtubules with nanometer precision. Towards this end, we determined the diameter of the helical trajectories (distance between adjacent maxima and minima in the sideways distance) for all 3D sliding events with at least two complete helical turns. We obtained a mean value of 86.2 ± 4.4 nm (N = 94, Fig. 3A) yielding an extension of the Ncd motors of about 18 nm (diameter of template microtubule [25 nm] + 2 × radius of transport microtubule [25 nm] + 2 × *in-situ* extension of Ncd motors [2 × 18] = diameter of the helical path [86 nm]; Fig. 3C). The diameter of the helical path showed no correlation with neither helical pitch nor sliding velocity (Supplementary Fig. 3). Another estimate for the *in-situ* extension of Ncd motors could be obtained from the sideways distance of sliding events that remained on the extreme right-hand side of surface-immobilized template microtubules (45.5 ± 0.1 nm; N = 83; Fig. 3B). This value yields an extension of the Ncd motors of about 21 nm (radius of template microtubule [12.5 nm] + radius of transport microtubule [12.5 nm] + *in-situ* extension of Ncd motors [21] = extreme right-hand sideways distance [46 nm]; Fig. 3C). Therefore, from our experimental measurements we estimated the *in-situ* extension of Ncd motors between crosslinked microtubules to be about 18 - 21 nm.

**Figure 3:**
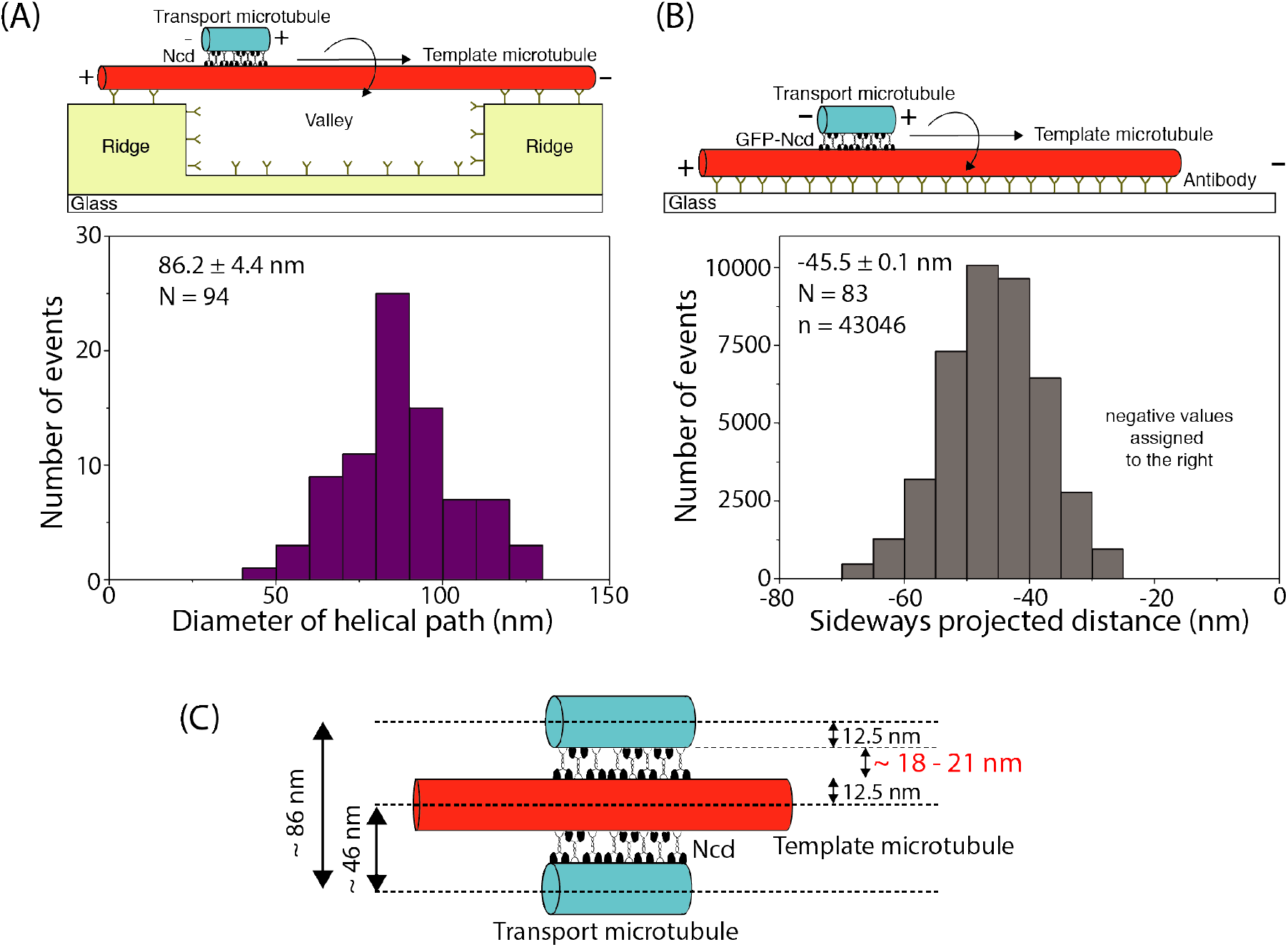
Spatial extension of Ncd motors between sliding microtubules. **(A)** Histogram corresponding to the diameter of the helical paths taken by antiparallel transport microtubules rotating around suspended template microtubules (3D sliding motility assay). The average diameter was 86.2 ± 4.4 nm (N = 94). The mean diameter of the helical paths was calculated from the peak-to-peak sideways distances obtained from the trajectories of antiparallel transport microtubules. **(B)** Histogram corresponding to the sideways distance of antiparallel transport microtubules sliding along the right-hand side of surface-immobilized template microtubule (2D sliding motility assay). The average sideways distance was −45.5 ± 0.1 nm (N = 83 events; n = 43046 data points). **(C)** As shown in the illustration, considering the diameter of the template and transport microtubule (25 nm) the *in-situ* extension of Ncd motors was calculated to be about 18 nm and 21 nm for the 3D and 2D sliding motility assays, respectively.

To elucidate the origin of the variation in the helical pitches of transport microtubules on suspended template microtubules (see Fig. 2A and Supplementary Fig.s 1A and B) beyond the sliding velocity and the microtubule length (Supplementary Fig.s 1C and D) we tested for the influence of the Ncd motor density. Unfortunately, direct measurements in our 3D sliding motility assays on suspended template microtubules were hampered for the following reasons: (i) It was not possible to reliably quantify the motor density from the intensity of the GFP signal of the motors due to the autofluorescence of the microfabricated polymer structures. (ii) Lowering the Ncd concentration in solution (to 0.4 nM) caused most transport microtubules to engage in an erratic helical motion (with only few events exhibiting complete rotations) even though the longitudinal motion along the template microtubules axis was not impeded. (iii) Increasing the Ncd concentration in solution (to 40 nM) led to unspecific motor binding to the polymer ridges causing the template microtubules to detach. We therefore turned to microtubule rotation measurements (i.e. the investigation of the rotational motion of microtubules around their own axes) in 2D motility assays on unstructured surfaces. First, we performed gliding motility assays on reflective silicon wafers, where rhodamine-speckled microtubules driven by Ncd motors (attached to the surface via Fab fragments and Anti-GFP antibodies) were imaged using fluorescence interference contrast (FLIC) microscopy ^15^. The recorded intensities of the speckles fluctuated periodically due to changes in speckle height with respect to the reflective surface, indicative of microtubule rotations (Fig. 4A). We observed that the rotational pitch and the sliding velocity of the microtubules reduced upon increasing the surface density of Ncd motors (Fig. 4B and Supplementary Fig. 4A). Secondly, we performed FLIC-based sliding motility assays on silicon wafers with rhodamine-speckled transport microtubules (Fig. 4C). While the helical motion of sliding transport microtubules around the surface-immobilized template microtubules was mostly blocked (as expected, see also Supplementary Fig. 2), we observed that the transport microtubules still rotated robustly around their own axes (Fig. 4C). At low Ncd concentration in solution (0.6 nM) only few transport microtubules rotated while at higher Ncd concentrations (2.4 nM and 6 nM Ncd) almost all transport microtubules rotated robustly. This behavior was confirmed when the Ncd concentration in solution was changed from 0.6 nM to 6 nM *in situ* during imaging. Transport microtubules which had not rotated periodically at 0.6 nM Ncd started to rotate robustly with short rotational pitches after switching to 6 nM Ncd (Supplementary Fig. 4E). Further, we observed that the rotational pitch and sliding velocity of the transport microtubules reduced upon increasing the Ncd concentration (Fig. 4D and Supplementary Fig. 4B). In summary, the rotational pitch of Ncd-driven gliding and sliding microtubules appears to be influenced by motor density. Interestingly, the *rotational pitches* of microtubules in these 2D gliding and sliding motility assays (0.5 - 2 μm) were in the same range as the *helical pitches* of microtubules in 3D sliding motility assays using suspended template microtubules.

**Figure 4:**
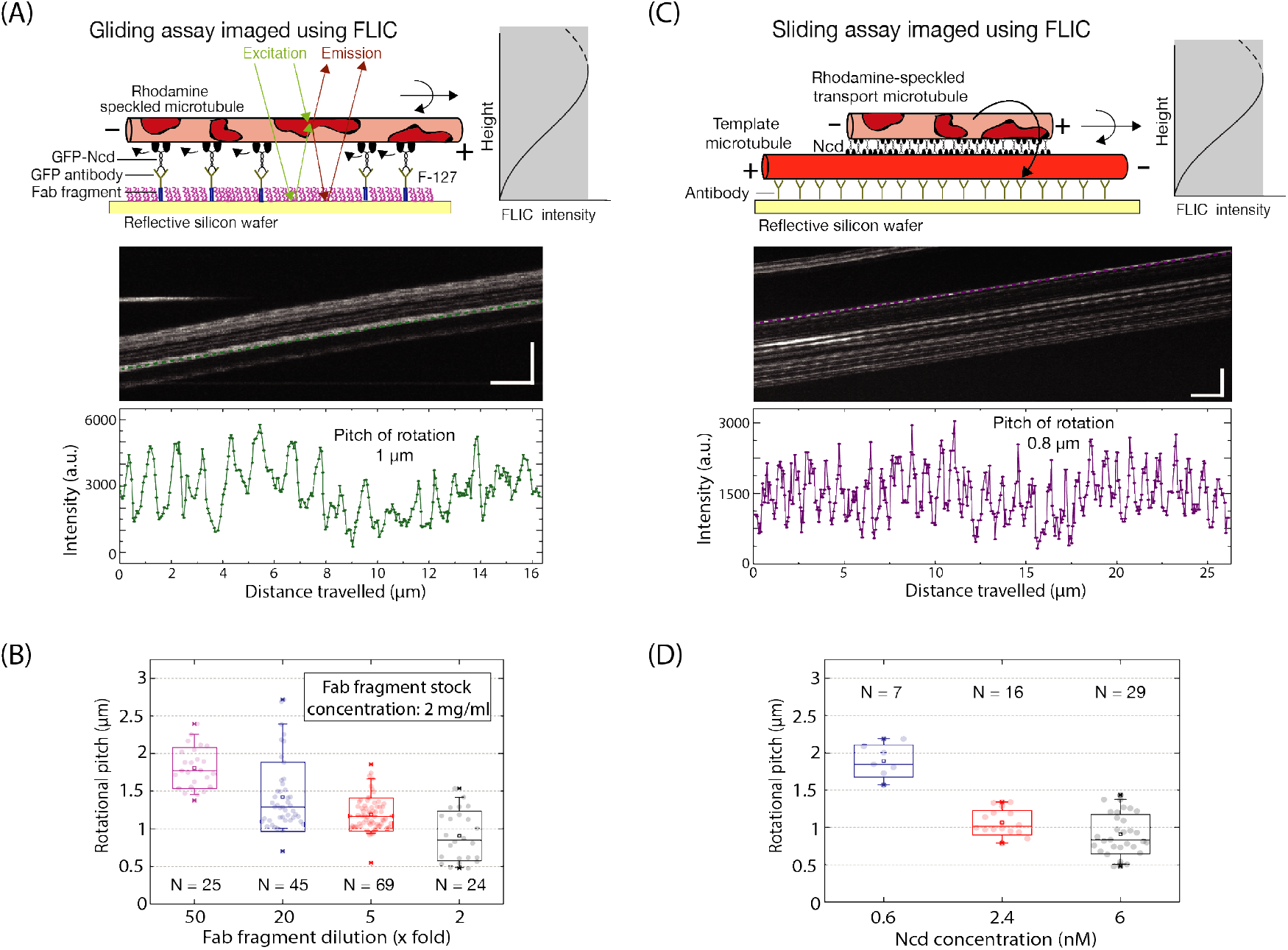
Dependence of rotational pitch of microtubules on Ncd motor density. **(A)** Measurement of the rotational pitch of speckled microtubules gliding on a reflective silicon substrate coated with GFP-Ncd via anti-GFP antibodies bound to Fab fragments. Due to fluorescence interference contrast (FLIC), the recorded intensities of the asymmetric speckles change as a function of height above the substrate. Rotational information of gliding speckled microtubules is interpreted from the periodic variation in the recorded intensity of the speckles on the microtubule (vertical scale bar: 10 μm; horizontal scale bar: 40 s in the example kymograph, FLIC intensity profile over time for one of the speckles indicated by the green line in the kymograph). The rotational pitch of this gliding microtubule is about 1 μm. **(B)** The rotational pitch of the gliding microtubules was dependent on the surface density of motors. The motor density was set by changing the Fab fragment dilution (Fab fragment stock concentration of 2 mg/ml; diluted 50x, 20x, 5x, and 2x) and ranged between 0.5-2.0 μm. **(C)** FLIC-based measurement of the rotational pitch of speckled transport microtubules sliding on surface-immobilized template microtubules (vertical scale bar: 10 μm; horizontal scale bar: 40 s in the example kymograph, FLIC intensity profile over time for one of the speckles indicated by the purple line in the kymograph). The rotational pitch of this sliding transport microtubule is about 0.8 μm. **(D)** The rotational pitch of the sliding transport microtubules was dependent on the motor concentration in solution (0.6 nM, 2.4 nM and 6 nM) and ranged between 0.5-2 μm.

## Discussion

Longitudinal motion of motors and cargo along surface-immobilized microtubules has been studied extensively in 2D *in vitro* motility assays. However, recent studies revealed that, in order to explore the complete 3D motion, the microtubules need to be suspended ^7,13,16,17^. This also became evident in our experiments where individual transport microtubules could freely move around the suspended parts of template microtubules, but were locked on the right-hand side of the surface-immobilized parts of the template microtubules (Fig. 2B, Supplementary Fig. 2B). Because locking occurred always on the right-hand side of the template microtubules, we reason that the helical motion on suspended microtubules is righthanded. The median helical pitch of 1.6 ± 0.2 μm cannot be related to the supertwist of the microtubules (all grown in the presence of GMPCPP and stabilized by taxol). The vast majority (about 96%^18^) of these microtubules are expected to comprise 14 protofilaments, which provide a left-handed supertwist of about 8 μm ^19,20^. Conceivably, the origin of the righthanded helical motion is related to the off-axis component in the power stroke of Ncd, elucidated in previous studies exploring the rotational motion of microtubules gliding on surface-bound Ncd motors ^8,9^. For parallel transport microtubules, the motors facing opposite directions between the crosslinked microtubules antagonize each other, locking the microtubules longitudinally as well as axially (Fig. 2C).

Based on our observations in FLIC measurements (Fig. 4B) we reason that transport microtubules exhibit right-handed rotations around their own axis in addition to their righthanded helical motion around suspended template microtubules (Figs. 1 and 2, see also illustration in Fig. 5A). To explain such complex movement, we carefully explored the geometry of the microtubules in the different motility assays (see Fig.s 5B and C for illustrations of the transverse sections of the microtubules). In gliding motility assays (Fig. 5B), the asymmetric power stroke of Ncd results in a right-handed rotational motion of the gliding microtubule around its own axis, as also observed in previous studies ^8,9^. In sliding motility assays (Fig. 5C), Ncd motors are oriented with the motor domains facing either the transport microtubule or the template microtubule. Ncd motor domains facing the transport microtubule are geometrically similar to the Ncd motors in the gliding assay - expected to rotate the transport microtubule around its own axis in a right-handed manner. In contrast, Ncd motors with motor domains facing the template microtubule cannot rotate the template microtubule (as it is fixed to the surface). Consequently, the off-axis power strokes of these motors are transposed to the transport microtubule, moving it helically around the template microtubule in a right-handed manner. Therefore, the two populations of Ncd motors enable the transport microtubules to undergo a combination of rotational motion around their own axes and helical motion around the template microtubules, both motions being right-handed with pitches in a similar range (0.5 – 2 μm).

**Figure 5:**
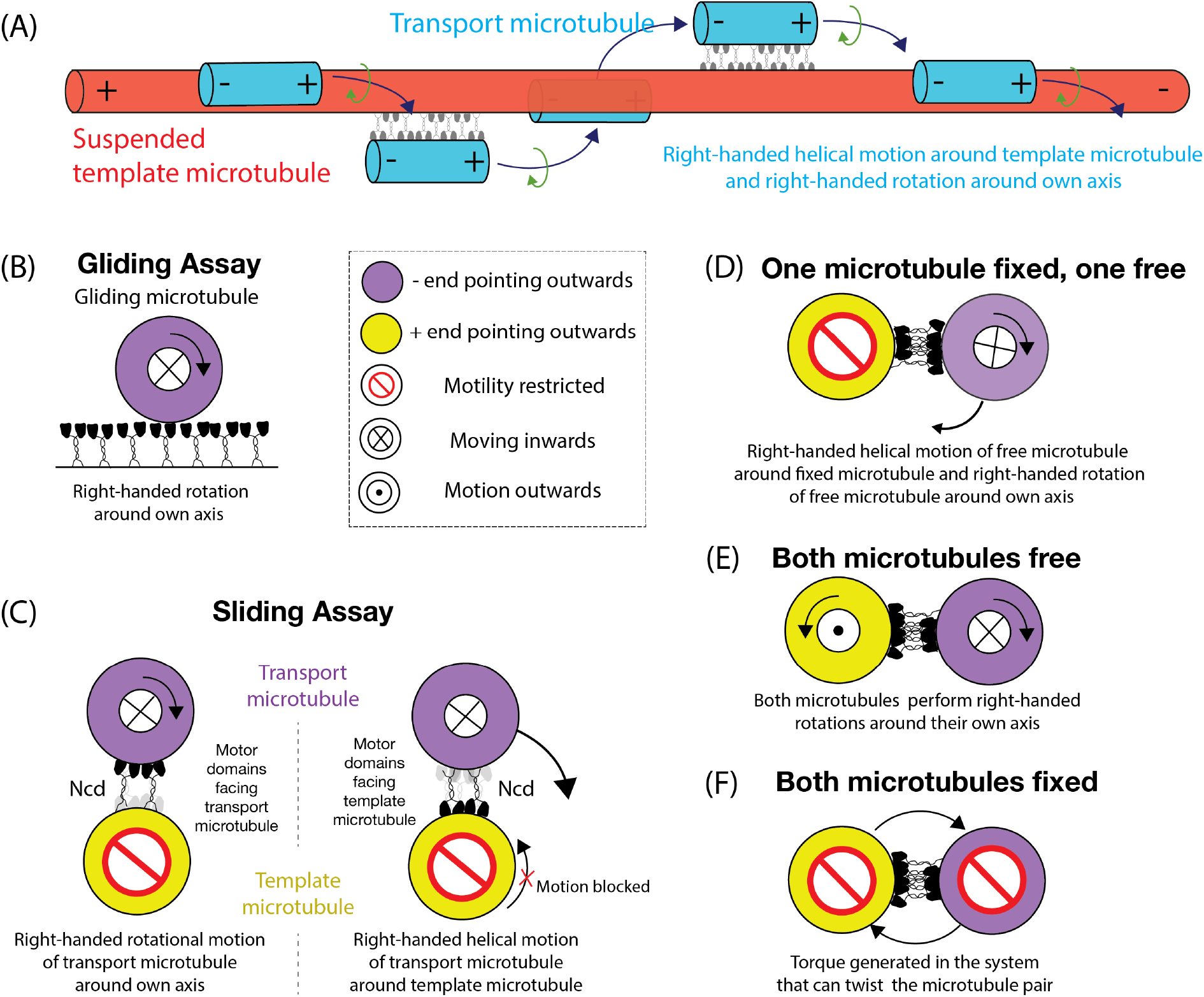
Illustrations to visualize the motion of antiparallel microtubules crosslinked by Ncd. **(A)** On a suspended template microtubule, antiparallel transport microtubules propelled by Ncd motors exhibit a right-handed helical motion around the template microtubules and a right-handed rotation around their own axes. **(B-C)** Transverse sections of microtubules propelled by Ncd in the geometries explored in this work. (B) In gliding motility assays, microtubules rotate in a right-handed manner while gliding forward. (C) In sliding motility assays, the Ncd motors bound via their tails to the template microtubule rotate the transport microtubule around its own axis (right-handed), similar to the gliding motility assays. The Ncd motors bound via their tails to the transport microtubule cannot rotate the surface-immobilized template microtubule. This blocked motion is transposed to the transport microtubule, which helically moves around the template microtubule in a right-handed manner. **(D-F)** Transverse sections of potential rotational motion and torque generation by microtubules crosslinked by Ncd *in vivo*. (D) If one microtubule is fixed somewhere and one microtubule is free, the free microtubule will move similar to what is observed for transport microtubules in our sliding motility assays on suspended template microtubules. The free microtubule will exhibit a right-handed helical motion around the template microtubule and a right-handed rotation around its own axis. (E) If both microtubules are free, they will only rotate around their own axis while sliding. (F) If both microtubules are fixed, they will coil around each other or twist their shape depending on how the microtubules are hinged and how strong the torque generated by the motors is.

The sizeable variation in the observed helical pitches and velocities of sliding transport microtubules cannot be entirely attributed to motor stochasticity as microtubule sliding involves multiple motors, a scenario where the stochasticity of single motors is supposedly averaged out. Also, we did not observe any correlation with the lengths of the transport microtubules, suggesting that the number of motors in the overlaps between template and transport microtubules alone does not influence helical pitch or velocity. Because it was difficult to systematically investigate the influence of the motor density in 3D sliding motility assays with suspended microtubules, we employed FLIC-based 2D gliding and sliding motility assays. We found that increasing the density of Ncd motors caused the microtubule velocity to decrease (Supplementary Figures 4A and B). This finding is in agreement with previous studies on kinesin-14 analogs (HSET ^21^ and XCTK2 ^22^) where a similar decrease in the sliding velocity was attributed to steric (longitudinal) hindrance between the motors ^21^. At the same time, we observed that the rotational frequency stayed rather constant upon variation of the motor density (Supplementary Figures 4A and B) with only slight impediment at very high motor densities. These findings suggest that the collective activity of Ncd motors influences the longitudinal motion to a larger extent than the rotational motion and provide a possible explanation for the resulting reduction in rotational pitch (defined as the ratio of sliding velocity divided by rotational frequency) with motor density. We thus reason that motor density, potentially varying between different microtubule overlaps in the same experiment, is the primary factor influencing the 3D trajectories of transport microtubules (both in terms of rotational and helical motion).

We speculate that Ncd-crosslinked, antiparallel microtubules can exhibit three geometries in the mitotic spindle *in vivo:* (i) One microtubules is fixed at some point (e.g. attached to the kinetochore^23^) while the other one is free (Fig. 5D). This geometry is similar to what we explore in our 3D sliding motility assays. The free microtubule would perform a right-handed helical motion towards the plus-end of the fixed microtubule, simultaneously rotating around its own axis. (ii) Both microtubules are free (Fig. 5E). In such a situation, both microtubules would perform right-handed rotations around their own axes, and would consequently roll around each other, while sliding apart. (iii) Both microtubules are fixed (e.g. in the stable microtubule overlap formed in the midzone of the mitotic spindle, Fig. 5F). In this geometry, the microtubules would coil around each other and twist the microtubule overlap. In a recent study it was shown that mitotic spindles in HeLa cells are indeed chiral, likely due to torques generated by kinesin-5 motor proteins^3^. It will be interesting to explore the magnitude of torque that kinesin-14, as well as other crosslinking kinesins (like kinesin-5^5,11^ and kinesin-8^6,24^), can exert. This can be made possible by extending our assay to enable torque measurements, for example by using 3D optical tweezers^13^. In a recent study^25^ it was shown that Ncd motors can only generate sub-pN forces along the longitudinal axis of the microtubule due to their diffusive tail anchorage. However, it is conceivable that in the axial direction the diffusivity of the Ncd tails is limited (i.e. the axial tail diffusivity between protofilaments is lower than the longitudinal tail diffusivity along a protofilament) and the motors might be able to generate significant torques.

In addition to resolving the 3D trajectories of microtubules sliding around each other, our experimental approach provides a compelling means to measure the extension of motors in their active state while crosslinking and sliding microtubules. For Ncd, we find an *in-situ* extension of 18 – 21 nm which is in the same range as estimated in earlier electron-microscopy studies for microtubule crosslinker mixtures in yeast cells, including the analog kinesin-14, klp2^26^. Interestingly, this distance is lower than estimated for passive crosslinkers from the MAP65 family, with protein extensions ranging between 25 – 35 nm^26–29^. This raises the intriguing question on what is the ‘real’ distance between crosslinked microtubules when crosslinked and sliding in the presence of multiple motor and non-motor crosslinkers. Is it possible that crosslinking motors exhibit a differential conformation, activity and interaction kinetics depending on how far they are extended? Provided that different microtubule crosslinkers have different natural extensions^26^, regulating the microtubule-microtubule spacing might be a smart means of controlling the activity of crosslinking motors and spatially sorting MAPs. Such sorting behavior has previously been reported for the actin bundling proteins, Fascin and a-Actinin^30^. Our approach is ideal to measure the extensions (and changes thereof) of different crosslinking motors and MAPs in their active states – in correlation with their functional behaviour as generators of longitudinal as well as axial motility and force.

In summary, this study elucidates that Ncd motors induce a helical motion of antiparallelly crosslinked microtubules around each other as well as a rotational motion around their own axis. *In vivo*, such motion (as opposed to a strict linear motion) might be useful to circumnavigate obstacles on microtubule lattices or in the surrounding environment and to redistribute MAPs. While minus-end directed kinesin-14 induces a right-handed rotational motion, other plus-end directed crosslinking kinesins, like kinesin-5, have been shown to induce left-handed rotational motion^5^. In microtubule overlaps where kinesin-5 and kinesin-14 co-exist, the two motors antagonize each other in the longitudinal direction ^22^ but the torques generated by them are expected to add up. Therefore, we hypothesize that in the midzone of mitotic spindles, where stable antiparallel microtubule overlaps are maintained for long periods, crosslinking kinesins build up torques in the system, thereby twisting and possibly coiling microtubules around each other. Finally, the mitotic spindle architecture appears to have an inherent right-handed (moving from spindle poles to center) chirality^4^. It is interesting to ponder if this chirality is just an evolutionary artifact that arises from the asymmetry in the powerstroke (or step cycle) of the evolutionarily conserved motor domains of kinesin motors or if there is any function associated with it.

## Methods

### Protein purification

Most experiments were performed with recombinant His_6_-tagged *Drosophila Melanogaster* full length GFP-Ncd expressed in SF9 insect cells and purified after cleavage of the His6-tag, as described previously^25^. For FLIC-based sliding motility assays, a different batch of full length GFP-Ncd expressed in *Escherichia coli* and purified as described previously (His6-tag not cleaved)^10^ was used.

### Fabrication of polymer structures on glass

Cleaned 22 × 22 mm^2^ glass coverslips (#1.5; Menzel, Braunschweig, Germany) were imprinted with a UV curable resin, EVG NIL UV/A 200nm (EV Group) using UV nanoimprint lithography (UV-NIL) as described in Mitra et al ^7^. The structure imprinted on the glass coverslips was characterized by repeated pattern of relief lines (that form the ridges) with a height of 370 nm (few experiments were also performed on 250 nm high ridges) and a width of 2 μm (or 5 μm), separated by 10 μm (the valleys between the ridges).

### Microtubule preparation

Both, template and transport microtubules, were guanylyl-(α,β)-methylene-diphosphonate (GMP-CPP) grown, taxol-stabilized (referred to as double stabilized). Template microtubules were long (average length > 15 μm) and rhodamine-labeled while transport microtubules were short (average length 1-2 μm) and Atto647n-labeled. In detail, template microtubules were grown in two cycles of polymerization. 4.6 μM rhodamine-labeled *porcine* tubulin was added to a polymerization solution, comprising of BRB80 (80 mM Pipes at pH 6.9, 1 mM MgCl_2_, 1 mM EGTA) supplemented with 1 mM GMP-CPP [Jena Bioscience, Jena, Germany] and 4 mM MgCl_2_, incubated on ice for 5 min followed by 30 min at 37 °C, to grow short microtubule seeds. The solution was centrifuged at 13300 rpm for 15 min at 25°C to remove free tubulin and the pellet was resuspended in a new polymerization solution supplemented with 0.4 μM of rhodamine-labeled tubulin. This solution was incubated overnight at 37°C with low tubulin concentration allowing microtubule seeds to anneal and form long microtubules. The solution was then centrifuged at 13300 rpm for 15 min at 25°C and the pellet was resuspended in BRB80T solution (BRB80 supplemented with 10 μM Taxol). Transport microtubules were grown as short Atto647n-labeled microtubule seeds in the same way as described for the first cycle of polymerization for template microtubules. For the FLIC-based sliding motility assays, Cy5-labeled template microtubules (grown as described above) and rhodamine-speckled transport microtubule (grown as described in Mitra et al.^15^) were used. For the FLIC-based gliding motility assays, rhodamine-speckled microtubules were used.

### Motility assays

Motility buffer (MB), used in all experiments, consisted of 20 mM Hepes at pH 7.2, 1 mM EGTA, 2 mM MgCl_2_, 75 mM KCl, 10 μM Taxol, 200 μg ml^−1^ casein, 10 mM dithiothreitol, 0.1% Tween-20, 20 mM D-glucose, 100 μg ml^−1^ glucose oxidase, 10 μg ml^−1^ catalase and either 1 mM ATP (MB-ATP) or 1 mM ADP (MB-ATP).

#### Ncd-driven microtubule sliding motility assays

3D sliding motility assays on suspended template microtubule were performed in microfluidic flow cells constructed on 22 × 22 mm^2^ glass coverslips patterned with UV-NIL polymer resin and 18 ×18 mm^2^ unpatterned glass coverslips, both dichlorodimethylsilane (DDS)-coated to make the surface hydrophobic, as described previously ^31^. Before silanization, the patterned coverslips were cleaned mildly (using 5 % mucasol and then in 70% ethanol) to avoid corrosion of the structure. 2D sliding motility assays on surface-immobilized microtubules were performed on unpatterned silanized coverslips or silicon wafers (10 × 10 mm^2^) with a 30 nm thermally grown oxide layer (for FLIC based motility assays; GESIM, Grosserkmannsdorf, Germany). Flow cells were flushed with the following sequence of solutions: (i) Bead solution consisting of 2% 200 nm Tetraspeck beads (incubation time 1 min). (ii) Antibody solution consisting of 20 - 100 μg ml^−1^ anti-rhodamine antibody (Mouse monoclonal clone 5G5; ThermoFisher Scientific) in PBS for unspecific binding of antibodies to the surface (incubation time 1 min). (iii) 1% pluronic F-127 in PBS (Sigma) in order to block the surface from unspecific protein adsorption (incubation time > 60 min). (iv) BRB80 washing step to remove unbound F-127 and exchange buffers. (v) Rhodamine-labeled template microtubule solution in BRB80T, followed by an immediate washing step with MB, in order to immobilize microtubules perpendicular to the ridges. (vi) MB-ADP solution containing Ncd (concentration ranging between 0.02 – 40 nM) for ADP bound motors to bind to template microtubules. (vii) MB-ADP solution containing Atto647n-labeled transport microtubules, followed by immediate washing step with MB-ADP, in order to crosslink few transport microtubules to the template microtubules and wash away the unbound ones. (viii) MB-ATP solution at the microscope after finding a suitable field of view. For FLIC-based sliding motility assays, Cy5-labeled template microtubules were immobilized on the surface using Anti-Cy5 antibodies (Mouse monoclonal clone CY5-15; Sigma-Aldrich) and rhodamine-speckled transport microtubules were used.

#### Ncd-driven microtubule gliding motility assays

For 2D FLIC-based gliding motility assays on antibody-coated surfaces (for Figure 4A), the assay was performed in flow cells constructed from DDS-coated silicon wafers and glass coverslips as described in Mitra et al.^15^ with replacement of BRB80 based motility buffer with MB-ATP. For Ncd-driven gliding motility assays on casein-coated surfaces (for Supplementary Figure 2), detergent cleaned (sonicated in 5 % mucasol and then in 70% ethanol) coverslips were used to construct the flow cells. Flow cells were flushed with the following sequence of solutions: (i) Casein buffer (20 mM Hepes at pH 7.2, 1 mM EGTA, 2 mM MgCl_2_, 75 mM KCl, 0.5 mg ml^−1^ casein) to passivate the surface with casein. (ii) MB-ATP solution containing Ncd (concentration ranging between 4-400 nM) for unspecifically binding Ncd motors on the surface via their tails (incubation time 5 min). (iii) MB-ATP solution containing rhodamine labeled microtubules to bind the microtubules to the motor coated surface (incubation time 1 min). (iv) MB-ATP solution to wash away unbound microtubules.

### Image acquisition

Optical imaging was performed using an inverted fluorescence microscope (Axio Observer Z1; Carl Zeiss Microscopy GmbH) with a 63x oil immersion 1.46NA objective (Zeiss) in combination with an EMCDD camera (iXon Ultra; Andor Technology) controlled by Metamorph (Molecular Devices Corporation). A LED white light lamp (Sola Light Engine; Lumencor) in combination with a TRITC filterset (ex 520/35, em 585/40, dc 532: all Chroma Technology Corp.) and an Atto647n filterset (ex 628/40, em 692/40, dc 635; all Chroma Technology Corp.), corresponding to rhodamine-labeled microtubules and Atto647n/Cy5-labeled microtubules respectively, were used for epifluorescence imaging. The imaging temperature was maintained at 24 °C by fitting a custom-made hollow brass ring around the body of the objective and connecting it to a water bath with a cooling/heating unit (F-25-MC Refrigerated/Heating Circulator; JULABO GmbH)^32^. For sliding motility assays, a field of view was selected when there were three (or more) Tetraspeck beads bound to the surface and several trackable template microtubules suspended between ridges. Template microtubules were imaged in the TRITC channel (for 50-100 frames at 3-10 fps with exposure time 100-300 ms) before and after imaging the transport microtubules in the Atto647n channel (for 5-15 min at 3-10 fps with exposure time 100-300 ms) to confirm that the template microtubules do not move while imaging the transport microtubules. For FLIC-based motility assays, a 63x water immersion 1.2NA objective (Zeiss) was used (higher working distance) in order to image rhodamine-speckled microtubule (for 5-15 min at 1 fps in the TRITC channel fps with exposure time 400 ms) on the silicon wafer surfaces on the far side of the flow cells.

### Image analysis

#### Sideways distance measurements

The acquired image streams of immobilized template microtubules (TRITC channel) and sliding transport microtubules (Atto647n channel) were analyzed using FIESTA ^14^. First, the beads, that serve as fiducial markers, were tracked in both the channels to obtain the color offset correction (nonreflective similarity: allowing for translation, rotation and scaling) between the two channels as well as the image drift correction in the corresponding channels. Template microtubules with relevant sliding events were tracked. Since template microtubules are immobilized, the filament position (tracked over 50-100 frames) was averaged to get an averaged filament position. Sliding transport microtubules (anti-parallel events) and locked transport microtubules (parallel events) on template microtubules with sliding events (to know the microtubule orientation) were tracked. After color and drift correction, the sideways distance was obtained as the perpendicular distance of the center point of the tracked transport microtubule from the averaged center line (tracked over the recorded 50-100 frames) corresponding to the template microtubule (illustrated in Figure 1B). For displaying the sideways distance plots, data was smoothened (often with the raw data in the background) by rolling frame averaging over 20 consecutive frames.

#### Analysis of helical sliding events on suspended microtubules

Rotational pitch, end-to-end velocity and the diameter of the helical path corresponding to each rotation of a helical sliding event was determined by manual computer-aided measurement of the sideways distance versus forward distance plots. For a given sliding event, measurements from individual rotations were averaged to obtain the mean pitch, velocity and diameter of helical path. Accounting for the helicity of the path traversed by the sliding microtubule, the actual velocity along the path (referred to as contour velocity in Supplementary Figure 1) was calculated assuming a helical path with a diameter of 86 nm 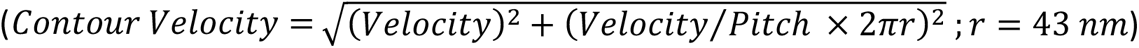.

#### *Rotational data analysis in the FLIC* motility *assays*

The rotational pitch of the gliding/sliding microtubules in the FLIC-based motility assays was obtained from their kymographs, which were generated in Fiji ^33^ using the MultiKymograph Plugin. The kymographs were then analyzed using the speckle analysis method described previously ^15^.

#### Analysis of distributions with error estimation

For estimating parameters from any given distribution (e.g. sideways distance, pitch, diameter of helical path) we used a bootstrapping approach^34,35^. Here, the distribution (N number of measurements) was resampled by randomly picking N measurements from the measured distribution (with replacement) and calculating the median of the resampled distribution. This was repeated 1000 times. The resulting bootstrapping distribution was used to estimate the parameter (mean of the bootstrapping distribution μ) and its error (standard deviation of the bootstrapping distribution σ). All values and errors as well as error bars in this paper use μ ± 3σ (99% confidence interval), unless otherwise noted.

## Supplemental Information

Supplemental Information consists four figures and three movies.

## Author Contributions

Author contributions: A.M., F.R., and S.D. designed research; A.M., R.G., R.R. performed research; F.R. contributed new reagents/analytic tools; A.M., R.G. analyzed data; and A.M. and S.D. wrote the paper with comments from the other authors.

## Acknowledgements

We thank Laura Meißner, Andrej Vilfan and Bert Nitzsche for scientific discussions and comments on the manuscript, all members of the Diez laboratory for fruitful interactions, Salvatore Girardo and the Microstructuring Facility at the Center for Molecular and Cellular Bioengineering at TU Dresden for preparing the polymer-structured coverslips and Corina Bräuer for technical support. We acknowledge financial support from the Deutsche Forschungsgemeinschaft through the Sonderforschungsbereich 1027 (Project A8), the Max Planck Institute for Molecular Cell Biology and Genetics Dresden and the Technische Universität Dresden.

---

**Supplementary Figure 1:**
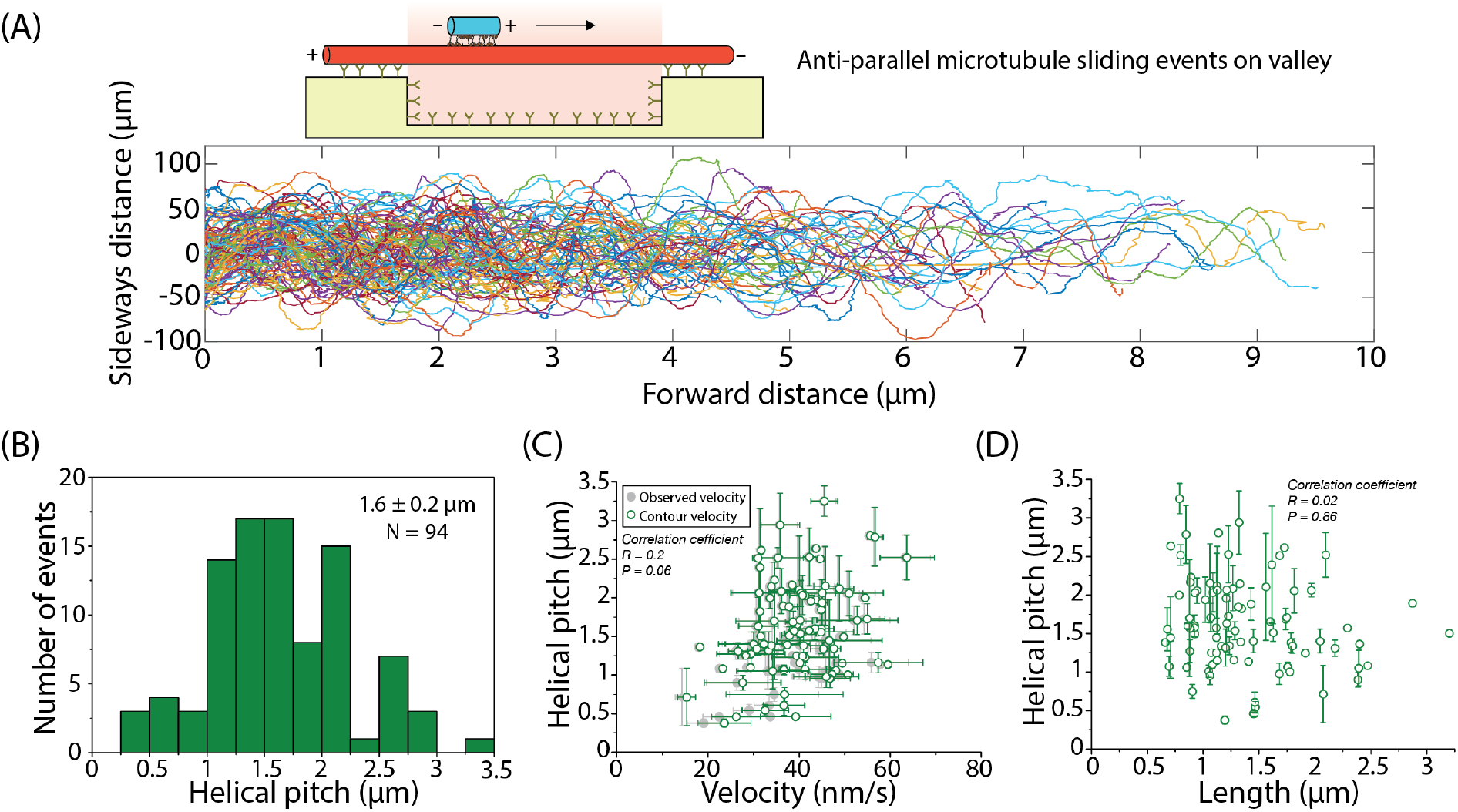
Transport microtubules exhibit variable helical pitches, independent of sliding velocity and microtubule length. **(A)** Sideways distance (rolling-frame averaged over 20 frames) of 94 antiparallel transport microtubules moving along the suspended parts of template microtubules. Each transport microtubule exhibited a robust helical motion with fairly constant helical pitch. Between various transport microtubules, helical pitches varied between 0.5 μm and 3 μm. **(B)** Histogram of helical pitches showing a median helical pitch of 1.6 ± 0.2 μm. **(C)** Helical pitch plotted with respect to the contour velocity of the sliding events. The contour velocity was calculated from the observed velocity (also plotted in the background in grey) accounting for the fact that the transport microtubules moved along a helical path around template microtubules. The actual (contour) velocity is therefore slightly higher than the observed velocity. There is no evident dependence of the helical pitch on the sliding velocity (R = 0.2, P = 0.06; Pearson correlation coefficient). **(D)** Helical pitch plotted with respect to the length of the sliding template microtubules. Smaller template microtubules showed slightly larger variations in helical pitch but no correlation with microtubule length was observed (R = 0.02, P = 0.86).

**Supplementary Figure 2:**
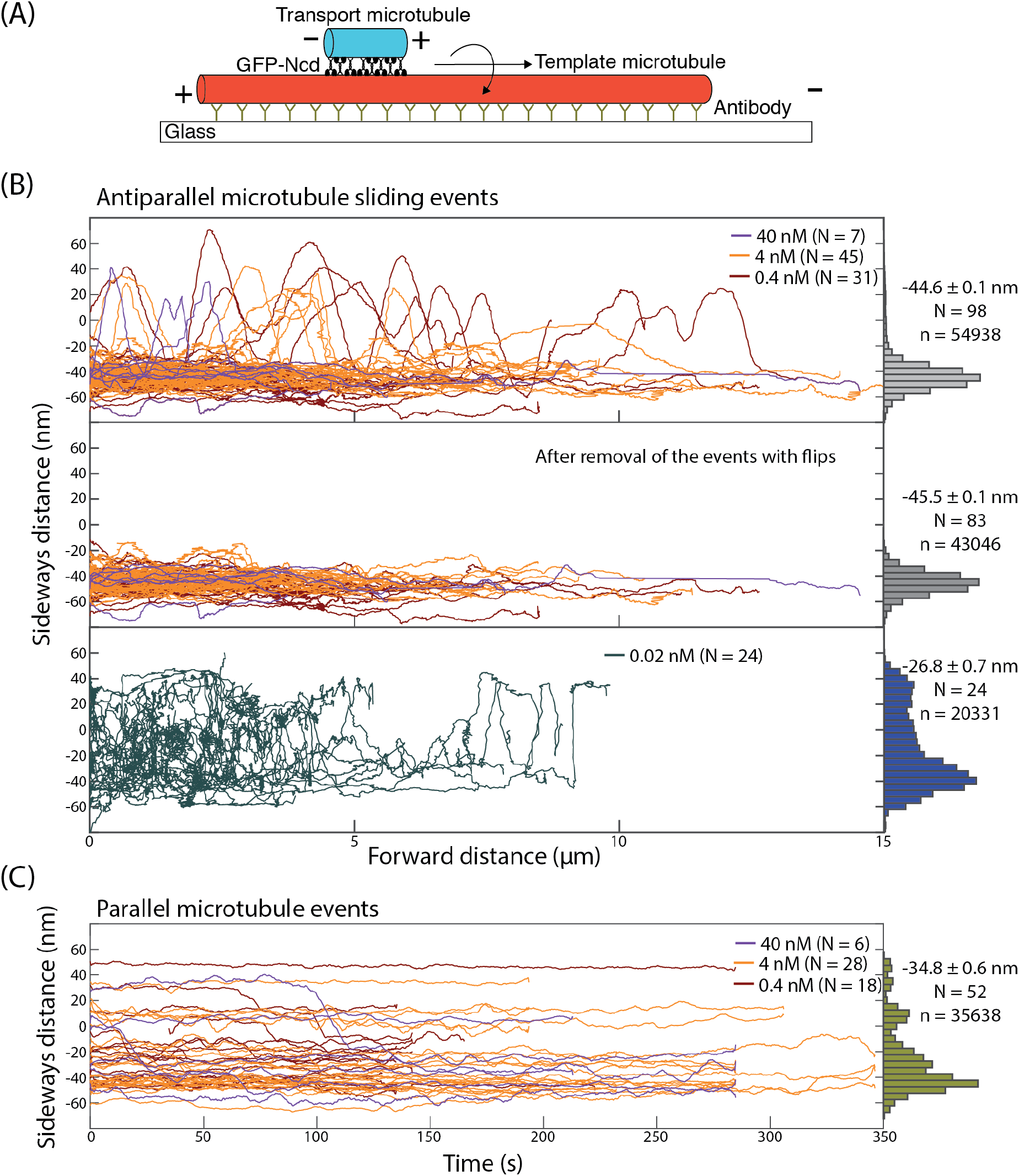
The 3D motion of transport microtubules is hindered in 2D sliding motility assays with surface-immobilized template microtubules. **(A)** Schematic representation of Ncd-driven sliding of an Atto647n-labeled transport microtubule along a surface-immobilized, rhodamine-labeled, template microtubule. **(B)** Sideways distance of antiparallel transport microtubules at different Ncd motor concentrations. At 0.4 nM, 4 nM and 40 nM Ncd motor concentrations (upper panel), most transport microtubules slid on the right-hand side of the template microtubule (see example sliding event at 4 nM Ncd motor concentration in Supplementary Movie 2). Some transport microtubules (15 out of 98) squeezed underneath the template microtubules and performed complete flips. The mean sideways distance was - 44.6 ± 0.1 nm (N = 98). When considering only the sliding events which remained on the right-hand side of the template microtubule, the average sideways distance was −45.5 ± 0.1 nm (N = 83). At 0.02 nM Ncd motor concentration the transport microtubules no longer remained on the right-hand side of the template microtubule but moved around erratically (in axial direction). There was still a slight preference for the right-hand side as indicated by the average sideways distance (−26.8 ± 0.7 nm; N = 24). **(C)** Sideways distance of parallel transport microtubules at different Ncd motor concentrations (0.4 nM, 4 nM and 40 nM). All events were locked, both in the longitudinal and axial direction (mean sideways distance of −34.8 ± 0.6 nm, N = 52). In all panels the displayed trajectories were smoothened (rolling frame averaged over 20 frames).

**Supplementary Figure 3:**
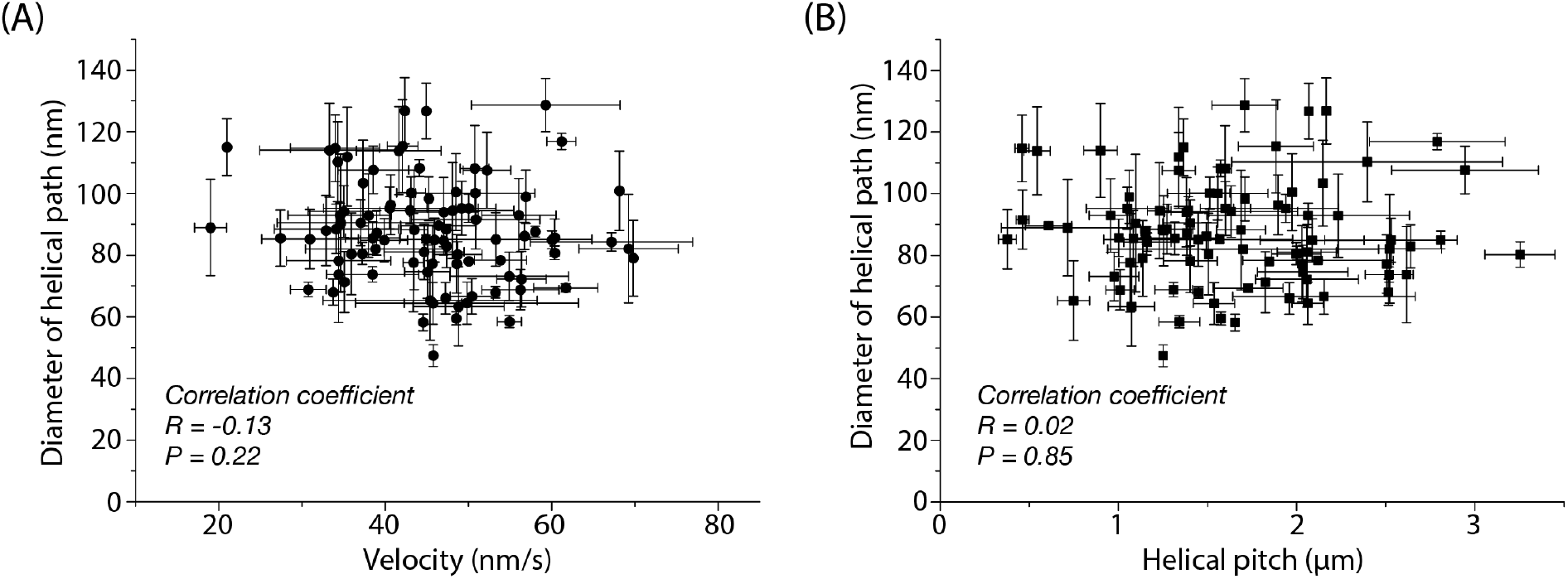
The spatial extension of Ncd motors between sliding microtubules is independent of sliding velocity and helical pitch. **(A)** Diameter of the helical path taken by antiparallel transport microtubules around suspended template microtubules plotted in dependence of sliding velocity. There was no correlation of the *in-situ* extension of Ncd motors with the microtubule sliding velocity (R = −0.13, P = 0.22; Pearson correlation coefficient). **(B)** Diameter of the helical path taken by antiparallel transport microtubules around suspended template microtubules plotted in dependence of the helical pitch. There was no correlation of the *in-situ* extension of Ncd motors with the helical pitch (R = 0.02, P = 0.85; Pearson correlation coefficient).

**Supplementary Figure 4:**
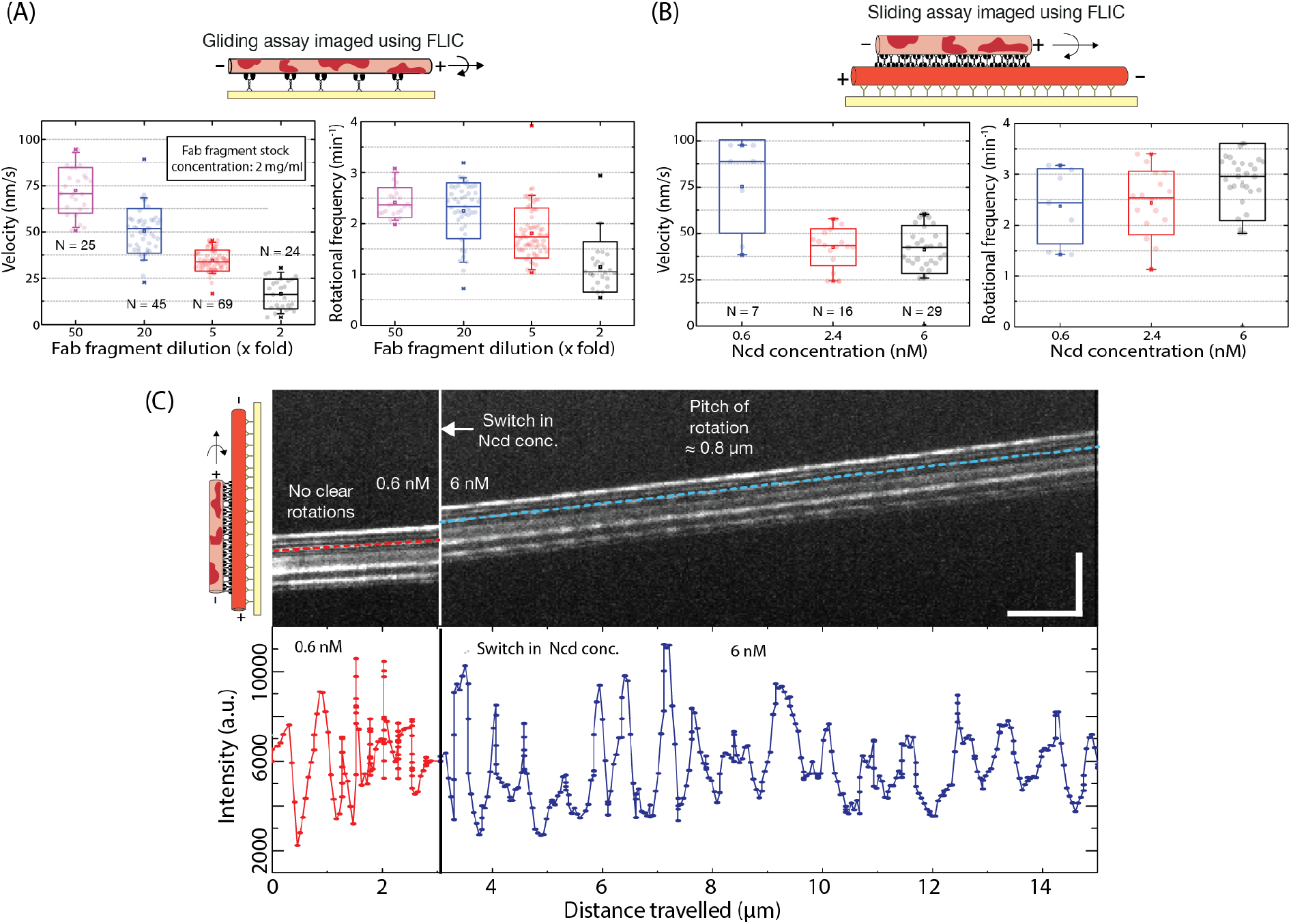
Details on FLIC-based gliding and sliding motility assays. **(A)** Box plots showing that gliding velocity and rotational frequency of microtubules gliding on Ncd motors are dependent on motor density. The rotational frequency saturates at 2 - 2.5 rotations per minute at low motor densities. The motor concentration was set by changing the Fab fragment concentration (stock concentration 2 mg/ml; diluted 50x, 20x, 5x and 2x). **(B)** Box plots showing that sliding velocity and rotational frequency of transport microtubules sliding on surface-immobilized template microtubules were slightly dependent on the motor concentration in solution. The rotational frequency ranged between 2.5 - 3 rotations per minute. **(C)** Kymographs and FLIC intensity profiles over time for one of the speckles indicated in the kymograph when the concentration of Ncd motors in solution was switched from 0.6 nM to 6 nM during imaging (horizontal scale bar: 40 s; vertical scale bar: 10 μm). At 0.6 nM Ncd the microtubule did not show any clear rotational motion. However, upon switching to 6 nM Ncd the microtubule rotated robustly around its own axis with a rotational pitch of about 0.8 μm.

**Supplementary Movie 1:** Example Atto647n-labeled transport microtubule (cyan) sliding along a rhodamine-labeled template microtubule (white) suspended over two valleys (10 μm wide) with a ridge (2 μm wide) in between, at 4 nM Ncd concentration in solution. The averaged tracked position of the template microtubule is indicated by the red line and the trajectory corresponding to the transport microtubule is indicated by the blue line. Details of this event are also shown in Fig. 1C.

**Supplementary Movie 2:** Example Atto647n-labeled transport microtubule (cyan) sliding along a rhodamine-labeled template microtubule (white) immobilized on a glass coverslip, at 4 nM Ncd concentration in solution. The averaged tracked position of the template microtubule is indicated by the red line and the trajectory corresponding to the transport microtubule is indicated by the blue line.

**Supplementary Movie 3:** Example rhodamine-speckled transport microtubule (cyan) sliding along a Cy5-labeled template microtubule (red) immobilized on a reflective silicon wafer, at 6 nM Ncd concentration in solution. The kymograph corresponding to this event is shown in Fig. 4C.

## Bibliography

1. Glotzer, M. The 3Ms of central spindle assembly: Microtubules, motors and MAPs. Nature Reviews Molecular Cell Biology (2009).

2. Cross, R. a. & McAinsh, A. Prime movers: the mechanochemistry of mitotic kinesins. Nat. Rev. Mol. Cell Biol. 15, 257–271 (2014).

3. Novak, M. et al. The mitotic spindle is chiral due to torques within microtubule bundles. Nat. Commun. 9, 3571 (2018).

4. Tolić, I. M. et al. Helical Twist and Rotational Forces in the Mitotic Spindle. Biomolecules 9, 132 (2019).

5. Yajima, J., Mizutani, K. & Nishizaka, T. A torque component present in mitotic kinesin Eg5 revealed by three-dimensional tracking. Nat. Struct. Mol. Biol. 15, 1119–21 (2008).

6. Bormuth, V. et al. The highly processive kinesin-8, Kip3, switches microtubule protofilaments with a bias toward the left. Biophys. J. 103, (2012).

7. Mitra, A., Ruhnow, F., Girardo, S. & Diez, S. Directionally biased sidestepping of Kip3/kinesin-8 is regulated by ATP waiting time and motor-microtubule interaction strength. Proc. Natl. Acad. Sci. U. S. A. 115, E7950–E7959 (2018).

8. Walker, R. A., Salmon, E. D. & Endow, S. A. The Drosophila claret segregation protein is a minus-end directed motor molecule. Nature 347, 780–2 (1990).

9. Nitzsche, B. et al. Working stroke of the kinesin-14, ncd, comprises two substeps of different direction. Proc. Natl. Acad. Sci. 113, E6582–E6589 (2016).

10. Fink, G. et al. The mitotic kinesin-14 Ncd drives directional microtubule-microtubule sliding. Nat. Cell Biol. 11, 717–23 (2009).

11. Kapitein, L.C. et al. The bipolar mitotic kinesin Eg5 moves on both microtubules that it crosslinks. Nature 435, 114–118 (2005).

12. Su, X. et al. Microtubule-sliding activity of a kinesin-8 promotes spindle assembly and spindle-length control. Nat. Cell Biol. 15, 948–957 (2013).

13. Bugiel, M., Mitra, A., Girardo, S., Diez, S. & Schäffer, E. Measuring Microtubule Supertwist and Defects by Three-Dimensional-Force-Clamp Tracking of Single Kinesin-1 Motors. Nano Lett. 18, 1290–1295 (2018).

14. Ruhnow, F., Zwicker, D. & Diez, S. Tracking single particles and elongated filaments with nanometer precision. Biophys. J. 100, 2820–8 (2011).

15. Mitra, A., Ruhnow, F., Nitzsche, B. & Diez, S. Impact-Free Measurement of Microtubule Rotations on Kinesin and Cytoplasmic-Dynein Coated Surfaces. PLoS One 10, e0136920 (2015).

16. Can, S., Dewitt, M. A. & Yildiz, A. Bidirectional helical motility of cytoplasmic dynein around microtubules. Elife e03205 (2014).

17. Brunnbauer, M. et al. Torque generation of kinesin motors is governed by the stability of the neck domain. Mol. Cell 46, 147–58 (2012).

18. Hyman, A. A., Chrétien, D., Arnal, I. & Wade, R. H. Structural changes accompanying GTP hydrolysis in microtubules: information from a slowly hydrolyzable analogue guanylyl-(alpha,beta)-methylene-diphosphonate. J. Cell Biol. 128, 117–25 (1995).

19. Nitzsche, B., Ruhnow, F. & Diez, S. Quantum-dot assisted characterization of microtubule rotations during cargo transport Accuracy of the 3-D quantum dot tracking. 1, 1–9

20. Ray, S. Kinesin follows the microtubule’s protofilament axis. J. Cell Biol. 121, 1083–1093 (1993).

21. Braun, M. et al. Changes in microtubule overlap length regulate kinesin-14-driven microtubule sliding. Nat. Chem. Biol. 13, 1245–1252 (2017).

22. Hentrich, C. & Surrey, T. Microtubule organization by the antagonistic mitotic motors kinesin-5 and kinesin-14. J. Cell Biol. 189, 465–80 (2010).

23. Redemann, S. et al. C. elegans chromosomes connect to centrosomes by anchoring into the spindle network. Nat. Commun. 8, 15288 (2017).

24. Su, X. et al. Mechanisms Underlying the Dual-Mode Regulation of Microtubule Dynamics by Kip3/Kinesin-8. Mol. Cell 43, 751–763 (2011).

25. Lüdecke, A., Seidel, A.-M., Braun, M., Lansky, Z. & Diez, S. Diffusive tail anchorage determines velocity and force produced by kinesin-14 between crosslinked microtubules. Nat. Commun. 9, 2214 (2018).

26. Roque, H., Ward, J. J., Murrells, L., Brunner, D. & Antony, C. The fission yeast XMAP215 homolog dis1p is involved in microtubule bundle organization. PLoS One (2010).

27. Gaillard, J. et al. Two Microtubule-associated Proteins of *Arabidopsis* MAP65s Promote Antiparallel Microtubule Bundling. Mol. Biol. Cell 19, 4534–4544 (2008).

28. Chan, J., Jensen, C. G., Jensen, L. C. W., Bush, M. & Lloyd, C. W. The 65-kDa carrot microtubule-associated protein forms regularly arranged filamentous cross-bridges between microtubules. Proc. Natl. Acad. Sci. 96, 14931–14936 (1999).

29. Subramanian, R. et al. Insights into Antiparallel Microtubule Crosslinking by PRC1, a Conserved Nonmotor Microtubule Binding Protein. Cell 142, 433–443 (2010).

30. Winkelman, J. D. et al. Fascin- and α-Actinin-Bundled Networks Contain Intrinsic Structural Features that Drive Protein Sorting. Curr. Biol. 26, 2697–2706 (2016).

31. Korten, T. et al. Fluorescence imaging of single Kinesin motors on immobilized microtubules. Methods Mol. Biol. 783, 121–37 (2011).

32. Ruhnow, F., Kloβ, L. & Diez, S. Challenges in Estimating the Motility Parameters of Single Processive Motor Proteins. Biophys. J. 113, 2433–2443 (2017).

33. Schindelin, J. et al. Fiji: An open-source platform for biological-image analysis. Nature Methods (2012).

34. Press, W. H., Teukolsky, S. a, Vetterling, W. T. & Flannery, B. P. Numerical recipes in C (2nd ed.): the art of scientific computing. Technometrics 29, (1992).

35. McCabe, G. P. & Moore, D. S. Introduction to the Practice of Statistics. (2006).

